# Precise insertion and guided editing of higher plant genomes using Cpf1 CRISPR nucleases

**DOI:** 10.1101/109983

**Authors:** Matthew B. Begemann, Benjamin N. Gray, Emma E. January, Gina C. Gordon, Yonghua He, Haijun Liu, Xingrong Wu, Thomas P. Brutnell, Todd C. Mockler, Mohammed Oufattole

## Introduction Paragraph

Precise genome editing of plants has the potential to reshape global agriculture through the targeted engineering of endogenous pathways or the introduction of new traits. To develop a CRISPR nuclease-based platform that would enable higher efficiencies of precise gene insertion or replacement, we screened the Cpf1 nucleases from *Francisella novicida* and Lachnospiraceae bacterium ND2006 for their capacity to induce targeted gene insertions via homology directed repair. Both nucleases, in the presence of guide RNA and repairing DNA template, were demonstrated to generate precise gene insertions as well as indel mutations at the target site in the rice genome. The frequency of targeted insertions for these Cpf1 nucleases, up to 8%, is higher than most other genome editing methods reported to date. Further refinements and broad adoption of the Cpf1 genome editing technology has the potential to make a dramatic impact on plant biotechnology.

## Main Body

Plant biotechnology using traditional molecular biology and transformation technologies has resulted in crops that have reduced energy intensive inputs, improved yields, and increased food, fuel, and fiber security across the globe. New advances in biotechnology, namely genome editing, have the potential to dramatically increase plant yields and bring product concepts to market that previously had large technical and/or regulatory barriers to market entry^1^. The adoption of genome editing in industrial plant biotechnology has resulted in new products with improved disease tolerance, yield, and food quality^2-4^. These advanced products were developed at a fraction of the time and cost of traditional ag-biotech traits. Broad adoption of this technology along with advances in genomics and breeding will result in innovation across the spectrum of native and novel traits including targeted allele swapping, trait stacking, and modification of expression elements of native genes^5^. The development of an easily accessible and functionally efficient genome editing platform will result in a new age of plant biotechnology.

Plant genomes can be edited using a variety of technology platforms including meganucleases, TALENs, CRISPR-Cas9, and zinc finger nucleases^6, 7^. Development of the CRISPR-Cas9 platform dramatically decreased the cost of performing genome editing experiments and allowed for expanded use of the technology in the plant sciences^8^. While easier to implement than TALENs, zinc fingers, and meganucleases, concerns with off target effects of CRISPR-Cas9 and its unclear intellectual property landscape have limited the broad adoption of CRISPR-Cas9 technology^9, 10^. Recently, an alternative family of CRISPR nucleases, Cpf1, has been identified and shown to function in editing the genome of human cells^11^. Cpf1 enzymes are a family of type V CRISPR nucleases that includes both endoribonuclease and endodeoxyribonuclease activities, allowing these nucleases to both process CRISPR-RNAs (crRNAs) and generate double strand breaks of DNA respectively^12^. The dual enzymatic activities allow for multiplex targeting from a single crRNA transcript^13^. In addition, Cpf1 nucleases have also been shown to have lower rates of off target edits relative to Cas9 nucleases^14, 15^.

Cpf1 nucleases in combination with crRNAs have been shown to generate indel mutations via non-homologous end joining repair (NHEJ) in both prokaryotic and eukaryotic systems^11, 16^. Recently, several of these Cpf1 nucleases were shown to generate indel mutations in both monocot and dicot plant species^17, 18^. While NHEJ mediated modifications have significant value for plant genome editing, enabling homology directed repair (HDR) modifications would expand the utility of this technology to even far more impactful applications such as the creation of superior alleles through site specific mutations and targeted insertions of genes and/or regulatory elements.

The Cpf1 nucleases from *Fransicella novicida* (FnCpf1) and Lachnospiraceae bacterium ND2006 (LbCpf1) were selected for investigation in plants with the goal of identifying CRISPR nuclease activities, when combined with the plant *in vivo* endogenous repairing events, that could produce gene knockouts and targeted insertions, both of which are inherently important to a robust plant genome editing platform. As originally described, LbCpf1 was able to generate indels in both *Escherichia coli* (*E. coli*) and human HEK293 cells, while the success of FnCpf1 was limited to *E. coli*^11^. Due to the inconsistencies of host specific performance of previously described genome editing nucleases^19^, it was not clear that negative or positive results in one system, whether *in vitro*, eukaryotic, or prokaryotic, would predict performance in another system. These inconsistencies are the result of the complex nature of heterologous expression of a genome editing system, which may result from suboptimal codon usage, the efficacy of nuclear localization tags, variable expression levels of each editing component, crRNA stability and other variables^20^. For these reasons, multiple Cpf1 nucleases were tested in our plant transformation system and assays performed to monitor both HDR and NHEJ repair genome edits.

To test the capability of each Cpf1 enzyme to generate targeted gene insertions via HDR, a screen was developed that would result in a visual phenotype upon genome editing. The Chlorophyllide-*a* oxygenase gene of rice (*CAO1*) was selected as a target due to an easily visualized “yellow” phenotype in homozygous plants, for loss-of-function of the *CAO1* gene^21^. In our experimental design, we included a hygromycin phosphotransferase gene (hpt), an antibiotic resistance marker, in the repair template (Fig. 1) that would both interrupt the rice *CAO1* coding sequence and expedite genetic screening. In each transformation experiment, rice (*Oryza sativa* cv. Kitaake) calli were bombarded with a set of three plasmids containing a Cpf1 expression cassette, a crRNA expression cassette specific to each Cpf1 enzyme, and a repair template with a hygromycin resistance gene under the control of the maize ubiquitin promoter flanked by 1kb regions of rice genomic DNA known as homology arms (Table S1-2). The Cpf1 genes were codon optimized for monocot plants, including an N-terminal nuclear localization tag, and were expressed by the 2x35S CaMV promoter. Two target sequences within exons of the *CAO1* gene were identified for target integration, both containing a TTTC PAM site, consistent with the PAM site requirements described previously for these enzymes^11^ (Fig. 1a). These sites are unique within the rice genome and were designed to minimized off target effects. Complete design of expression systems and repair templates can be found in Supplementary figures S1-S6.

**Figure 1:**
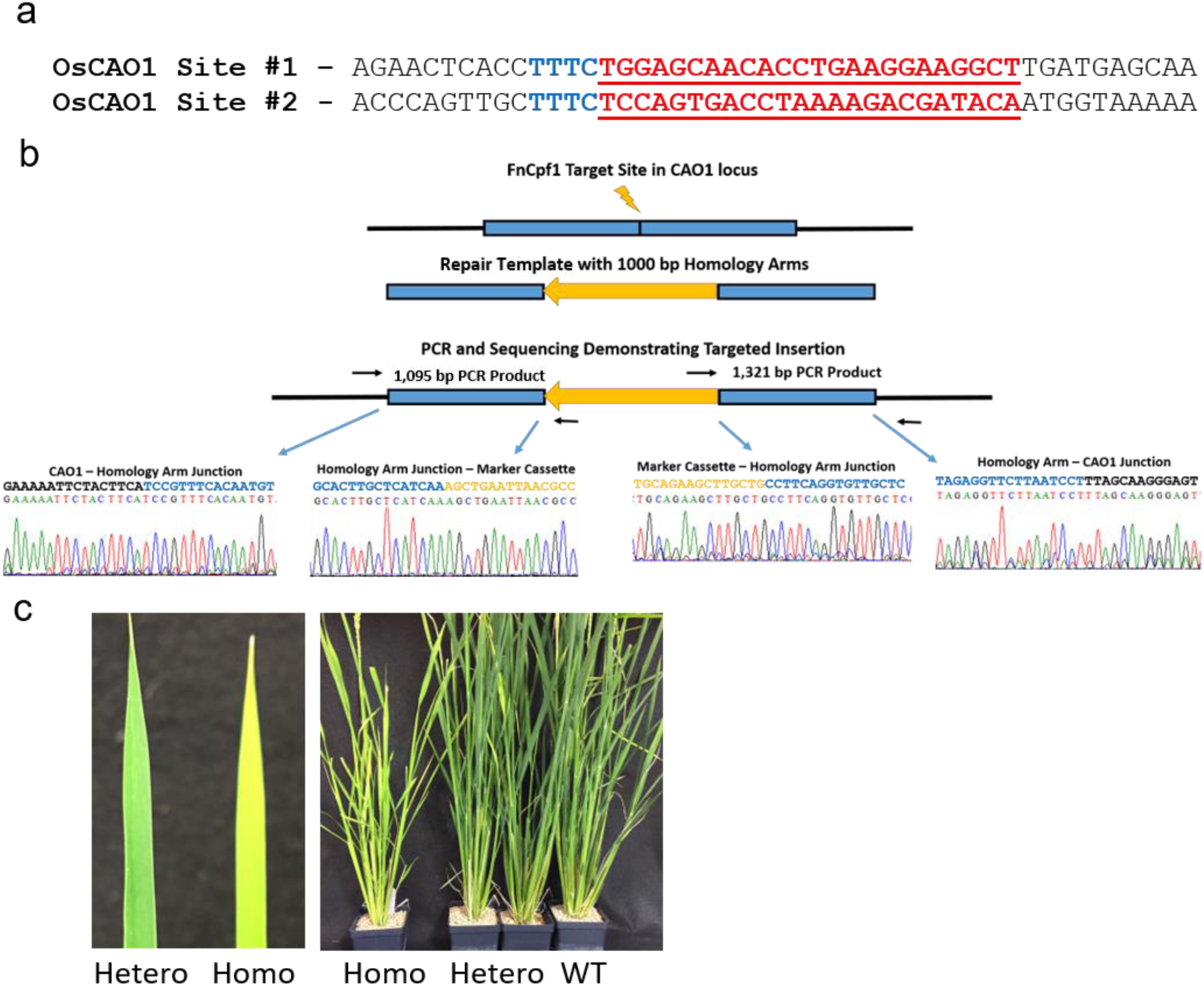
Targeted integrations of a hygromycin resistance marker into the *CAO1* gene of rice. (a) Target sites for crRNA design. Site #1 is shown as the reverse complement for clarity. The PAM site is identified in blue and the target sequence is in red. (b) Schematic and sequence alignment of hygromycin insertion (yellow sequence) at *CAO1* site #1 using FnCpf1. (c) Comparison of leaves and whole plants of homozygous *CAO1* lines and heterozygous or wild type lines.

A set of plasmids specific to FnCpf1 and *CAO1* site #1 (Experiment GE0001) were bombarded into rice calli and hygromycin resistant calli were identified. Fragments of resistant calli were screened via PCR for the presence of a targeted integration of the resistance marker into the *CAO1* gene via HDR. PCR assays were developed to amplify junction fragments from the genomic region outside the homology arm into the resistance marker (Fig. 1b). Amplification would only occur in the presence of a targeted integration event. Out of 36 independent lines, 3 generated PCR products for both the upstream and downstream junctions, for a targeted integration frequency of 8% (Fig. S7 and Table 1). PCR fragments from callus line number 6 (GE0001-6) were sequenced and aligned to the predicted insertion sequence (Fig. 1b). These alignments demonstrated a clean junction between the genomic DNA, homology arms, and both ends of the resistance marker. Sequencing of additional PCR fragments from independent callus lines GE0001-1 and GE0001-21 demonstrated a similar result of precise targeted integration events (Fig. S8). In brief, all independently identified targeted insertions resulted in precise genomic site integration events, leaving intact junction regions between genomic DNA and upstream homology arm DNA, and downstream homology arm and corresponding genomic DNA.

**Table 1:** Results of Rice Biolistics Experiments

| Experiment | Target | Nuclease | Resistant Callus | Insertions | Indels |
| --- | --- | --- | --- | --- | --- |
| GE0001 | OsCAO1 Site 1 | FnCpf1 | 36 | 3 (8%) | 1 (3%) |
| GE0001-R | OsCAO1 Site 1 | FnCpf1 | 96 | 8 (8%) | 31 (32%) |
| GE0031 | OsCAO1 Site 1 | LbCpf1 | 55 | 0 (0%) | 1 (2%) |
| GE0046 | OsCAO1 Site 2 | LbCpf1 | 96 | 3 (3%) | 10 (10%) |

Callus line GE0001-1 was placed on regeneration medium, resulting in four sibling plantlets (T0), with each plantlet PCR positive for both the upstream and downstream junction elements (Fig. S9). T1 seed were collected individually from each putative transgenic T0 plant. The T1 seeds from each transgenic plant were germinated and screened via qPCR for the presence of the hygromycin resistance marker. T1 seed from GE0001-1.3 were planted and approximately 25% of seedlings were pale green, indicative of 3:1 Mendelian segregation. Upon growth in the greenhouse, T1 plant GE0001-1.3.1 was found to be homozygous for insertion via qPCR. As expected, this line produced a yellow leaf phenotype (Fig. 1c) and is absent of Chlorophyll *b* (Fig. S10), consistent with a loss of function of the *CAO1* gene^21^.

To determine if FnCpf1 generated mutations outside of the target site, a preliminary analysis was performed to identify off target cutting. Cas-OFFinder^22^ was used to identify the top off target sites within the rice genome, both of which contained a 4 bp mismatch. Both genomic regions were amplified via PCR from multiple GE0001 events as well as from wild type rice. These fragments were subcloned, sequenced via Sanger sequencing and compared to the wild type sequences. No off-target mutations were detected at these genomic loci.

The FnCpf1 targeted integration experiment into *CAO1* site #1 (GE0001-R) was repeated on a larger scale of transformation screening to demonstrate the efficacy of the FnCpf1 nuclease to generate targeted gene insertions and the reproducibility of the system (Table 1). This experiment resulted in a similar frequency of targeted integrations (8%). This rate of targeted integrations is higher than other published experiments using the *S. pyogenes* Cas9 enzyme in plants^2, 23, 24^. These results are especially significant for early proof of concept work with limited optimization of delivery methods. With wider adoption and additional optimization, this genome editing platform has the potential to deliver even higher rates of targeted integrations, which would enable adoption of genome editing at all stages of a plant biotechnology program.

In addition to FnCpf1, LbCpf1 was also screened for its ability to generate targeted integrations via HDR. Biolistics experiments were performed by using LbCpf1 and respective crRNAs and repair templates for *CAO1* targets #1 and #2 (GE0031 and GE0046, respectively). Targeted integration events, as determined by PCR screening of callus tissue, were identified for both target sites at a frequency of 0% and 3%, respectively (Table 1). Sequencing of PCR amplicons of the upstream junction of events from GE0046 simply demonstrated an insertion that matches the design of the repair template (Fig. S11). These data demonstrate the ability of a second Cpf1 nuclease to generate targeted integrations via HDR in plants.

In addition to screening callus from the previously presented experiments for targeted gene insertions, these callus fragments were also screened for the presence of indels at the target site. While these experiments were initially designed to generate HDR, there was the expectation that indels generated by NHEJ should be present as well. Calli from each experiment were screened using the T7 Endonuclease I (T7EI) assay. Indels were identified via T7EI at both sites within the *CAO1* gene with frequencies ranging from 2-32% (Table 1 and Fig. S12-S13). PCR products from callus with a positive result from the T7EI assay were subcloned and sequenced (Fig. 2a and 2b). Indel sizes ranged from 3-75 bp (median 7 bp) for FnCpf1 and 3-15 bp (median 8 bp) for LbCpf1. Multiple T0 plants were regenerated from experiment GE0046 calli that were shown to be positive for indels at the callus screening stage. The target region of these plants was amplified, subcloned, and sequenced (Fig. S14). One of the T0 plants (GE0046-40) is clearly a chimera, while the other two are hemizygous or bi-allelic.

**Figure 2:**
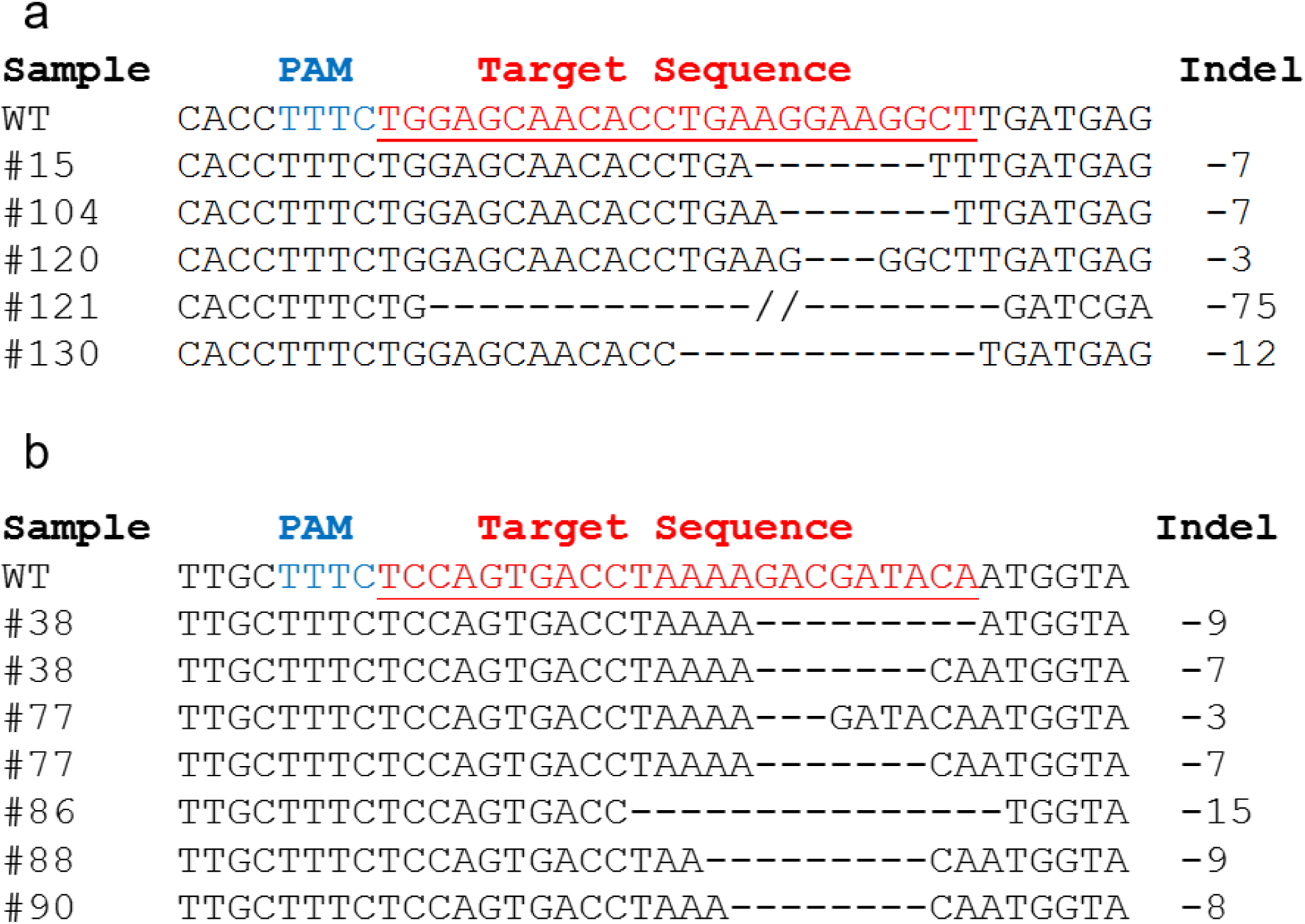
Identification of indels generated by FnCpf1 and LbCpf1 in the *CAO1* gene. (a) Alignment of indels identified from FnCpf1 experiments GE0001 and GE0001-R at *CAO1* site #1. (b) Alignment of indels identified from LbCpf1 experiment GE0046 at *CAO1* site #2.

The position of all identified indels are distal to the PAM site, which is consistent with the known enzymology of these nucleases^11^. The size and positioning of these mutations are similar to those recently published using Cpf1 nucleases in dicots and monocots^17, 18^. Our observations are also similar to those presented in mice and human cells^11, 16^. Callus from FnCpf1 appeared to have one predominant population of indel mutants, while samples from LbCpf1 tended to have multiple mutant populations. The observed frequencies of indel mutations in our biolistics experiments were lower than those observed with *Agrobacterium* mediated delivery of FnCpf1 and LbCpf1 in rice and tobacco^17, 18^. Clearly, optimization of component delivery has the potential to improve the frequency of edits. It is also interesting that FnCpf1, which has no to limited nuclease activity in human cells^11, 15^, performs especially well in plant systems, both in our experiments and other reports^17^. This again demonstrates the variability in the functionality of genome editing nucleases across hosts.

The results presented here demonstrate that both FnCpf1 and LbCpf1, when used together with crRNA and repairing template DNA, are capable of meditating both NHEJ and HDR based genome editing in plants. This is the first demonstration to our knowledge of Cpf1-mediated DNA insertion in any eukaryotic system. These results are significant with respect to both advancement of Cpf1-based genome editing technology and plant biotechnology. The demonstration of Cpf1 to generate targeted insertions will expand the application of this genome editing platform across basic and applied science disciplines. The frequencies of targeted insertions demonstrated that the platform we established by using these nucleases, with the potential to increase with additional optimization of repairing template DNA and reagent delivery methods, etc., will enable wide-scale adoption of this technology in plant biotechnology from trait discovery to product development and trait stacking.

## Online Methods

### Plasmids Construction

Cpf1 sequences were codon optimized for monocot plants and synthesized by Genscript. Cpf1 genes were subcloned via restriction digest cloning into expression vectors with a pUC19 backbone containing the 35S promoter and 3’UTR as well as a 35S terminator. Repair template plasmid was assembled by using the hot fusion cloning methodology^25^. Briefly, homology arms were amplified from rice genomic DNA and assembled with the hpt cassette driven by maize ubiquitin promoter and terminated by 35S in a pUC19 backbone crRNA sequences were synthesized as G-Blocks from IDT with the rice U6 RNA promoter and terminator.

### Transformation

Biolistic-mediated transformation of rice embryogenic calli derived from mature seeds of *Oryza sativa* L. cv. Kitaake was performed as described in Chen *et al*.,^26^ with slight modifications. DNA amounts of Cpf1, crRNA and repair template plasmids used in bombardment experiments were 0.5µg, 0.5µg and 1µg per single shot respectively. The premixed DNA constructs were co-precipitated onto 0.6-µm gold microprojectiles as previously described^2^. Callus samples were bombarded once using PDS 1000/He biolistic system (BIO-RAD) at helium pressure of 650 psi with a target distance of 6 cm and 27 in Hg vacuum in the chamber. Post-bombardment culture, hygromycin-B selection, and plant regeneration were performed as previously described^2^ with minor modifications.

### PCR Screening and Sequencing

DNA was extracted from rice callus and leaf tissue using the CTAB DNA extraction method. 100ng of purified genomic DNA was used for all PCR reactions. PCR assays for insertion screening were performed using GoTaq polymerase (Promega) and the primers listed in supplementary Table 2. PCR products were electophoresed on 1% agarose gels before visualization. PCR products were gel extracted and cloned into sequencing vectors using the pJET system (ThermoFisher). Plasmids were sequenced using Sanger sequencing by services of Genscript.

### T7EI Screening

Target regions were amplified by using the PCR method described above. Cleaned up PCR products were treated with T7 Endonuclease I (New England Biolabs) in a 20*µ*L reaction. Reactions were incubated at 37°C for 15min followed by the addition of 1.5*µ*L 0.25M EDTA to stop the reaction. Digests were electrophoresed on a 2% agarose gel prior to visualization. PCR products from calli positive for an indel were cloned and sequenced as described above.

